# Hepatic Transcriptomic Landscape of Chicken Reveals Host Responses to Spotty Liver Disease

**DOI:** 10.64898/2026.03.21.713329

**Authors:** Varsha Bommineni, Lekshmi K. Edison, Chaitanya Gottapu, Gary D. Butcher, Subhashinie Kariyawasam

## Abstract

Spotty Liver Disease (SLD) is an acute bacterial infection of layer chickens in production, caused by *Campylobacter hepaticus*, and occurs most frequently in barn-housed and free-range systems. The disease is characterized by a sharp decline in egg production and increased mortality. The hallmark pathological feature is 1-2 mm white to grey necrotic foci distributed across the liver surface. Despite its growing economic impact on commercial poultry, the molecular mechanisms underlying host responses to *C. hepaticus* infection remain poorly understood. To address this gap, we performed a comprehensive transcriptome analysis of liver tissue from chickens naturally infected with SLD compared to uninfected controls. High-throughput transcriptome sequencing, yielding 9,277 differentially expressed genes (DEG), of which 3,063 were upregulated and 6,214 were downregulated. Functional pathway enrichment analysis revealed significant alterations in immune and metabolic processes associated with SLD pathophysiology. Infected chickens exhibited significant activation of immune response pathways, particularly cytokine-cytokine receptor interactions involving interleukins *IL-22, IL-21,* and *IL-6,* along with enhanced cell signaling, and cell adhesion. Among the individual genes, *C1QTNF1* and the adhesion molecule gene *ADGRD1* were notably overexpressed, indicating enhanced inflammatory activity. In contrast, core hepatic metabolic functions were profoundly reduced, as evidenced by downregulation of oxidative phosphorylation, fatty acid metabolism, iron ion binding, and heme binding pathways. A marked increase in serum amyloid A gene (*SAA*) expression further confirmed robust acute-phase responses and compromised liver function during infection. Together, these findings demonstrate a complex interplay between inflammatory activation and metabolic dysregulation during SLD. The strong upregulation of acute-phase proteins and pro-inflammatory cytokines demonstrates the host’s vigorous attempt to combat bacterial infection, whereas the concurrent suppression of essential metabolic pathways reflects the pathological consequences of SLD. This study provides a transcriptomic characterization of host responses to *C. hepaticus* infection, offering insights into SLD pathogenesis and potential avenues for targeted intervention.

## 1. Introduction

Spotty Liver Disease (SLD) is an emerging bacterial disease of layer chickens in production that has become a major economic concern for the global egg industry. This disease is primarily caused by *Campylobacter hepaticus* (1,2), while *Campylobacter bilis* has been identified as a less common causative species (3). Outbreaks have been reported in the United States, United Kingdom, New Zealand, Jordan, and across Europe (4–6). Outbreaks typically occur during the peak laying period, between 22 and 35 weeks of age, mostly during warm conditions (7) and are associated with flock mortalities of up to 10% and production losses of 10–35% (8).The resulting reduction in egg production and survivability makes SLD one of the most important health challenges in layer operations.

SLD is characterized by the sudden death of affected hens, with a few birds showing prior clinical signs such as lethargy, pyrexia, and in some cases icterus (8). This disease is more prevalent in layers, which are more vulnerable than broilers or non-laying chickens, perhaps as a result of physiological or hormonal factors (7). The characteristic postmortem lesion of SLD is in the liver, where 1–2 mm grey-white lesions are distributed across the surface of the liver parenchyma (9). Histologically, these foci represent multifocal hepatocellular necrosis, often accompanied by fibrin deposition and infiltration of heterophils and other inflammatory cells (10). Notably, lesions are not restricted to specific hepatic structures but appear randomly distributed, suggesting a systemic process. Although such damage is expected to affect liver function, it is unclear how exactly it contributes to mortality and loss of production. These unresolved questions highlight the need for molecular approaches to uncover the host–pathogen interactions driving hepatic pathology (11). In chickens, the liver performs diverse functions, including detoxification, bile secretion, carbohydrate, protein and fat metabolism, vitamin storage, (12) and serves as a unique immune organ that contributes to pathogen defense. It eliminates pathogens through innate immune cells such as Kupffer cells, dendritic cells, and natural killer cells, while adaptive immunity is supported by resident T and B lymphocytes (13). Previous work on immune responses of SLD has demonstrated that infected hens develop protective antibodies and can resist reinfection, with seroprevalence studies revealing variable antibody responses (2-64%) across affected flocks (14). Limited studies suggest that *C. hepaticus* infection triggers proinflammatory responses, including elevated levels of IL-1β, IL-6, and IL-8 (15,16). However, the hepatic immune response, the key site of pathology in SLD, remains poorly characterized.

To bridge this knowledge gap, we conducted high-throughput RNA sequencing to analyze the liver transcriptomes of hens naturally infected with SLD and exhibiting lesions characteristic of the disease. We aimed to uncover key host responses and disrupted pathways associated with SLD pathogenesis. These findings contribute to a deeper understanding of liver-specific host–pathogen interactions and may highlight targets for improved diagnostics, prevention, and control strategies.

## 2. Materials and Methods

### 2.1. Sample Collection and Study Design

Infected livers were obtained from 31-week-old White Leghorn commercial layer chickens during an active SLD outbreak at a commercial egg production facility that was experiencing elevated mortality and decreased egg production and was free from other diseases. Postmortem examination of deceased birds revealed characteristic SLD lesions, including multiple white-to-tan foci distributed across the liver surface. The presence of *C. hepaticus* was confirmed through microbiological isolation and PCR-based identification of affected tissues (17). PCR assays also excluded the presence of *C. billis* (*18*). Negative control liver samples were collected from age-matched laying hens (31-week-old) sourced from a SLD-free commercial facility with no history of *C. hepaticus* infection. All liver samples were collected during necropsy procedures, immediately stabilized in RNAlater (Thermo Fisher Scientific, Lenexa, KS, USA) solution, transported on ice, and stored at −80°C until RNA extraction. For transcriptomic analysis, three independent biological replicates from each infected and control liver were selected for downstream processing. All samples were obtained as part of routine disease surveillance activities conducted under veterinary supervision.

### 2.2. RNA Extraction and rRNA Depletion

Total RNA was extracted from liver tissue samples using the RiboPure RNA Purification Kit (Invitrogen, Waltham, MA, USA) following the manufacturer’s protocol to ensure optimal yield and purity. Ribosomal RNA (rRNA) was depleted from the total RNA preparations using the RiboMinus Eukaryote Kit v2 (Invitrogen, Waltham, MA, USA) and the enriched messenger RNA (mRNA) fraction was concentrated using the RiboMinus Concentration Module (Invitrogen, Waltham, MA, USA) to achieve the optimal input concentration for high-throughput sequencing library preparation. RNA quality and quantity were determined at each step using a Qubit 4.0 Fluorometer (Thermo Fisher Scientific, Wilmington, DE, USA) to ensure that all samples required standards for downstream transcriptomic analysis.

### 2.3. RNA Sequencing Library Construction and Transcriptomic Analysis

Purified chicken liver mRNA (100 ng per sample) was processed for sequencing library construction using the TruSeq Stranded mRNA Library Prep Kit (Illumina In*C.*, San Diego, CA, USA). The enriched mRNA was enzymatically fragmented and used for double-stranded cDNA synthesis. The resulting cDNA fragments were subjected to 3′ end adenylation and ligated with TruSeq RNA Combinatorial Dual Index Adapters (Illumina In*C.*, San Diego, CA, USA), followed by PCR enrichment to amplify the indexed libraries. The prepared cDNA libraries were pooled, distributed across two flow-cell lanes, and sequenced on the Illumina NovaSeq X Series platform (Illumina In*C.*, San Diego, CA, USA). Raw sequencing data were subjected to quality control processing using CLC Genomics Workbench v.24.0.04 (Qiagen, Redwood City, CA, USA), including adapter trimming and removal of poor-quality reads (Phred score < 30). Quality-filtered reads were aligned to the White Leghorn layer chicken reference genome (NCBI

Ref Seq Assembly GCF_016700215.2 & Gen Bank Assembly GCA_016700215.2) for transcript quantification. Differential gene expression analysis between SLD-affected and control liver samples was performed using CLC Genomics Workbench v.24.0.04 to identify differentially expressed genes (DEGs). Statistical significance thresholds were established at p-value < 0.01, false discovery rate (FDR) < 0.05, and fold change (FC) > 2 (19,20). Each experimental group consisted of three independent biological replicates to ensure statistical robustness.

### 2.4 Quantitative Real-Time PCR Verification of RNA-seq Data

Total RNA from chicken liver samples was isolated using the RiboPure RNA Purification Kit (Invitrogen, Waltham, MA, USA) according to the manufacturer’s instructions. A total of 100 ng of RNA from liver control and infected liver samples was reverse transcribed into cDNA using the iScript cDNA Synthesis Kit (Bio-Rad Laboratories, Hercules, CA, USA). To validate the expression trends observed in the RNA-seq dataset, selected chicken genes were examined by reverse transcription quantitative PCR (RT-qPCR) with gene-specific primers listed in S1 File.

Reactions were prepared using SsoAdvanced Universal SYBR® Green Supermix (Bio-Rad Laboratories, Hercules, CA, USA) and run on a QuantStudio™ 5 Real-Time PCR System (Applied Biosystems, Carlsbad, CA, USA). The thermal cycling program included an initial denaturation at 95 °C for 3 minutes, followed by 40 cycles at 95 °C for 15 seconds and 60 °C for 30 seconds. Transcript levels were normalized against the housekeeping gene β-actin, which served as the internal control.

### 2.5 Data Availability

The transcriptomic profile data (raw and processed) described in this study were deposited in the Gene Expression Omnibus (GEO) database in NCBI under accession number GSE325026

## 3. Results

### 3.1 Transcriptomic Profiling Reveals Extensive Gene Expression Changes

Quality trimmed, high-throughput RNA sequencing reads were mapped to the White Leghorn chicken reference genome (NCBI RefSeq Assembly, GCF_016700215.2) with an average global mapping rate of 70.7% (range: 65.0-80.5%). After quality control and normalization, 25,462 genes were retained for differential gene expression analysis. The complete, unfiltered differential gene expression dataset are provided in detail in S2 File. The analysis identified 9,277 significantly DEGs in the liver samples, comprising 3,063 upregulated and 6,214 downregulated genes. The volcano plot illustrated the transcriptomic changes contributing to SLD pathogenesis, with significant DEGs showing fold changes in both directions (Fig 1A). Expression density plots showed apparent transcriptional differences between control and SLD-affected samples (Fig 1B), whereas a heatmap analysis demonstrated clear clustering of genes and expression pattern differences between infected and control livers (Fig 1C).

**Fig 1.**
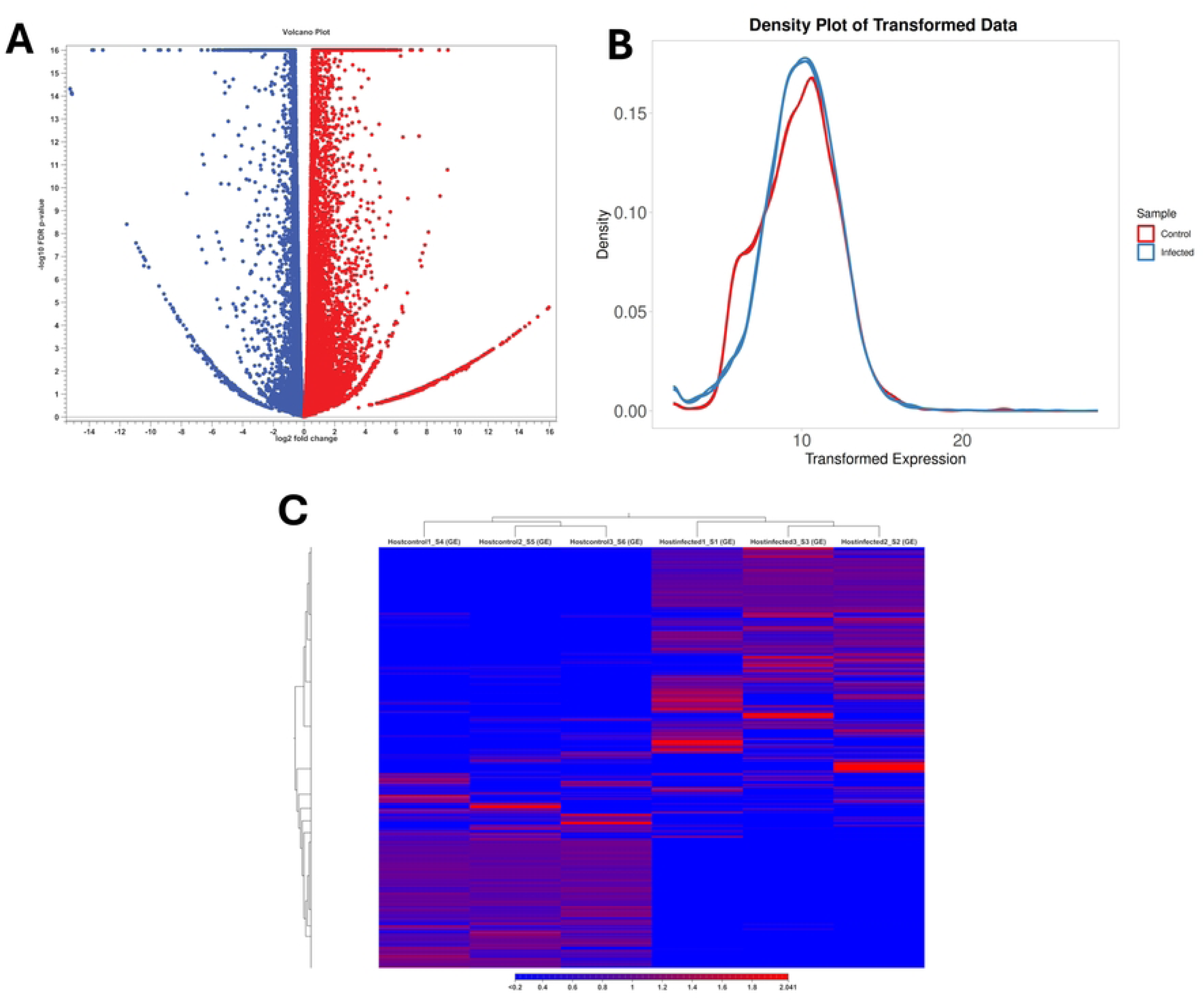
Transcriptomic analysis of *C. hepaticus* infection in chicken liver. (A) Volcano plot depicting the differential transcriptomic profile of *C. hepaticus* in infected chicken liver. In the volcano plot, the x-axis represents log₂ fold change and the y-axis represents –log₁₀ FDR p-value. Each dot represents a gene, with red indicating significantly upregulated genes and blue indicating significantly downregulated genes. (B) Density plot showing the distribution of normalized gene expression values in infected and control liver samples. (C) Hierarchical clustering heatmap of differentially expressed genes (DEGs) in infected and control chicken liver samples.

### 3.2 Gene Ontology (GO) Enrichment Analysis

GO enrichment analysis was performed using g:Profiler. GO terms were classified into three standard categories: Biological Process (BP), Molecular Function (MF), and Cellular Component (CC). For upregulated genes, g:Profiler identified 291 significantly enriched GO terms (49 MF, 196 BP, and 46 CC; provided in detail in S3 File as upregulated GO terms), whereas downregulated genes yielded 172 significant GO terms (33 MF, 89 BP, and 50 CC; provided in detail in S4 File as downregulated GO terms). To simplify interpretation, the top 30 GO terms in the downregulated and upregulated categories are presented in (Figs 2A and 2B), respectively.

**Fig 2.**
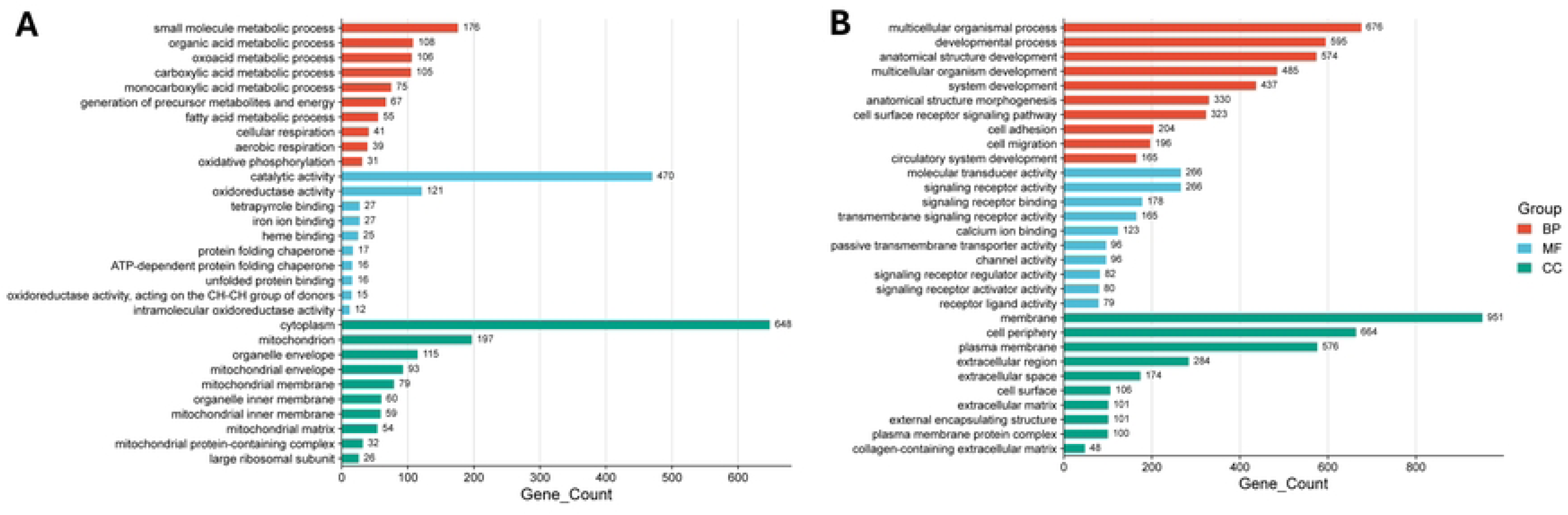
GO Enrichment analysis of differentially expressed genes in infected chicken liver. (A) downregulated genes and (B) upregulated genes. Bar plot showing the significantly enriched GO terms categorized under BP, MF, and CC. The x-axis represents the number of upregulated genes associated with each GO term (gene count), while the y-axis lists the GO terms. The color of each bar indicates the GO category

Statistical significance was determined using the Benjamini–Hochberg false discovery rate (FDR) with a threshold of p < 0.05. Upregulated GO pathways were predominantly enriched for processes related to cell surface signaling receptor activity, cell migration, cell adhesion, calcium ion binding, and developmental processes. In contrast, downregulated GO pathways were significantly associated with small molecule metabolic process, fatty acid metabolic process, Iron ion binding, heme binding, and oxidoreductase activity, consistent with broad suppression of core hepatic metabolic functions in hens with SLD.

### 3.3 Kyoto Encyclopedia of Genes and Genomes (KEGG) Analysis

KEGG pathway enrichment analysis (g:Profiler) identified nine and 20 significantly enriched pathways among the upregulated and downregulated DEGs, respectively; provided in detail in S2 File and S3 File as upregulated and downregulated KEGG terms. To highlight the most biologically relevant processes, the top pathways are shown in (Figs 3A and 3B). Upregulated pathways were dominated by cytokine–cytokine receptor interaction, cell adhesion molecules, extracellular matrix (ECM)–receptor interaction, and focal adhesion, consistent with immune activation and tissue remodeling. In contrast, downregulated pathways were primarily associated with metabolic pathways, oxidative phosphorylation, carbon metabolism, and fatty acid degradation, reflecting marked suppression of mitochondrial and core metabolic functions in SLD-affected hens.

**Fig 3.**
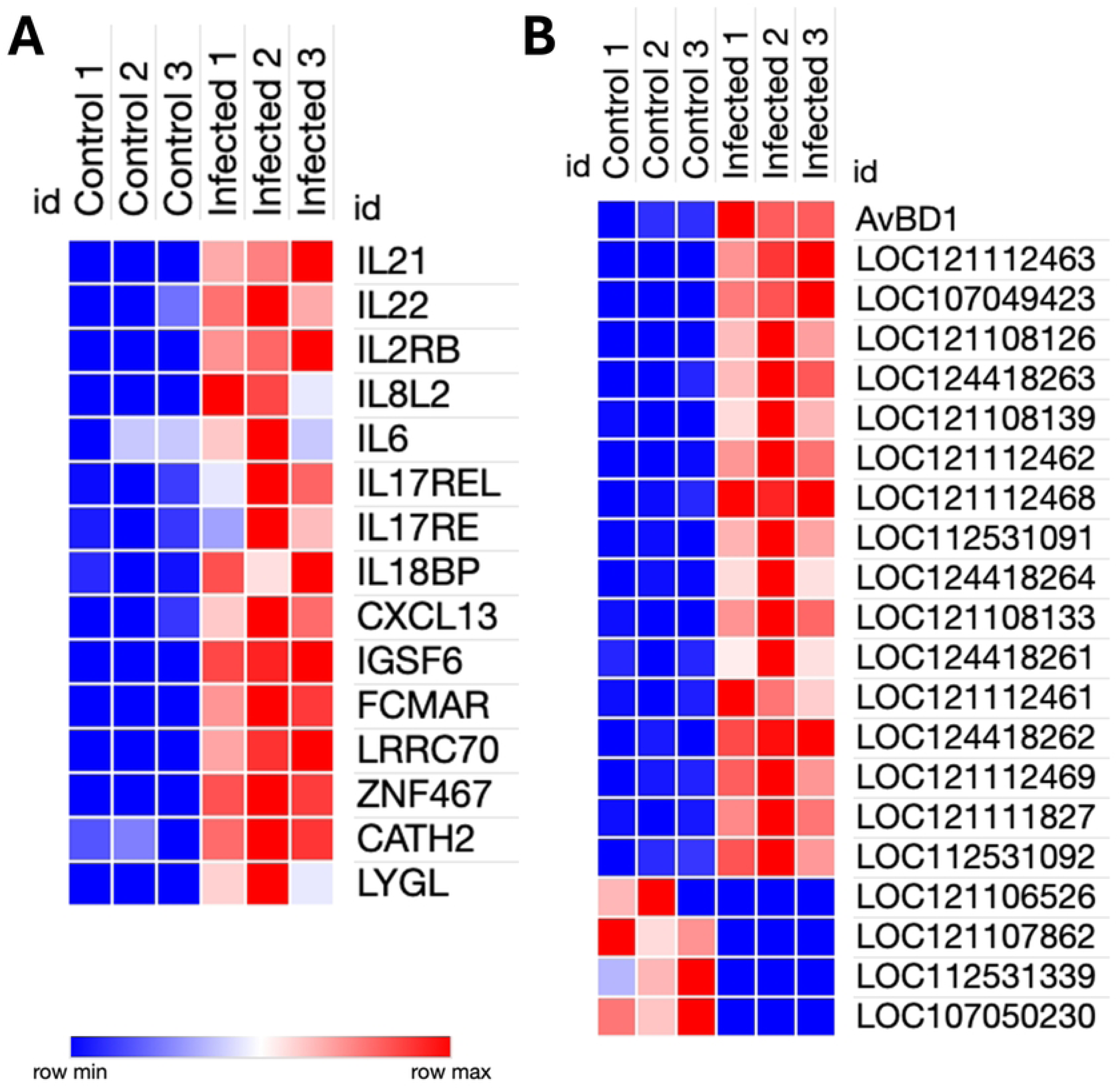
KEGG pathway enrichment analysis of differentially expressed genes in infected chicken liver. (A) Upregulated genes and (B) downregulated genes. The dot plot illustrates the top significantly enriched KEGG pathways derived from the differentially expressed genes. The x-axis represents the number of genes involved in each pathway, and the y-axis lists the pathway names. The color gradient corresponds to the −log₁₀(p-value), where darker red indicates higher statistical significance, and the dot size reflects the number of genes contributing to each pathway.

### 3.4 Patterns of Highly Expressed Gene Modules

#### Immune and inflammatory pathways

Elevated expressions were observed in immune mediator genes such as *IL2RB, IL21, IL22, IL8L2, IL-6, IL17REL, IL17RE, IL18BP* and the chemokine gene *CXCL13*, accompanied by up upregulation of multiple immunoglobulin variable region genes (*LOC121112463, LOC107049423, LOC121108126, LOC124418263, LOC121108139, LOC121112462, LOC121112468, LOC112531091, LOC124418264, LOC121108133, LOC124418261, LOC121112461, LOC124418262, LOC121112469, LOC121111827, LOC112531092)* suggesting B-cell activation. Increased expression was also noted in immune receptor and regulator genes, including *IGSF6, FCMAR, LRRC70, and ZNF467*. The *LYGL* (lysozyme G-like) was also upregulated, indicating the activation of innate antibacterial defense mechanisms.

Additionally, the avian beta-defensin (*AVBD1*) and *CATH2* genes were upregulated, indicating the activation of antimicrobial peptide defenses against bacterial infection. Together, these findings indicate robust innate inflammatory and humoral immune responses in SLD-affected livers, further supported by the marked upregulation of the serum amyloid A (*SAA*) gene. In contrast, transcripts associated with NK and T cell activation (e.g., *LOC121106526* and loci annotated to immune response-regulating pathways *LOC121107862, LOC112531339, LOC107050230*) were downregulated (Fig 4).

**Fig 4.**
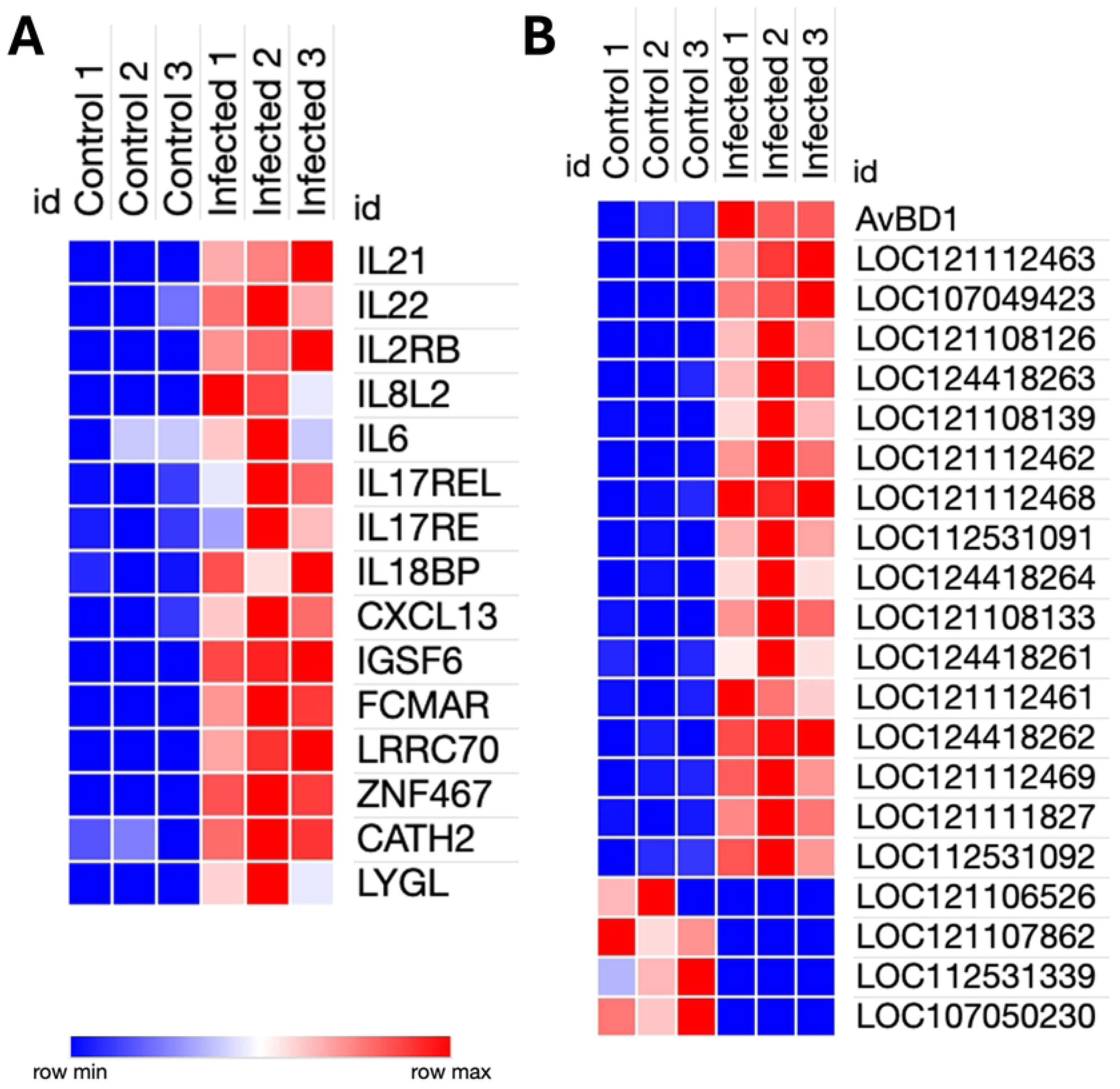
Heatmap of differentially expressed immune and inflammatory genes in control and infected samples. Infected1, Infected 2, and Infected 3 represent the three biological replicates of the liver-infected group; and Control1, Control 2, and Control 3 represent the three biological replicates of control groups.

#### Structural integrity

Genes related to the ECM and cytoskeletal structures were broadly suppressed, including collagen-associated genes (e.g., *LOXL4*, collagen alpha-1(I)-like gene), intermediate filaments (*KRT75*), junctophilin-2 like, outer dense fiber protein 3–like, and coiled-coil domain proteins, consistent with matrix degradation and cytoskeletal disruption. In contrast, *TUBB2A* (tubulin beta 2A class IIa), which is essential for microtubule-based cell migration, was upregulated. SKA1 (spindle and kinetochore-associated complex subunit 1), which regulates chromosome segregation and mitotic cell cycle progression, was also upregulated. Additionally, *PERPA* (periplakin A), a cell-cell adhesion protein, was upregulated, suggesting the activation of cellular adhesion mechanisms during infection.

#### Metabolic dysfunction

Core hepatic pathways were markedly downregulated, including those involved in carbohydrate metabolism (*FBP2, HKDC1, ALDOB, PDHA2, GLUL*), lipid metabolism (*APOB, APOA2, PLIN1*), amino acid and urea cycle processes (*OTC, PYCR1*), detoxification (*CYP2C18, CYP2AC1*), and mitochondrial/energy homeostasis and transport (*ACAA1, SLC30A2*), collectively indicating metabolic dysfunction. Several genes involved in iron homeostasis, heme biosynthesis, and iron-dependent metabolic processes (*FTH1, FECH, MELTF, ISCU, ACO2,* and *RRM2*) were also downregulated.

#### Cell signaling and ion transport

Signaling receptor genes *ACKR4* (chemokine scavenger) and *ADGRD1* (adhesion GPCR) were upregulated, along with ion/transport regulator genes (*ANO9, GABRA5, CAP2*), suggesting altered chemokine gradients, G protein-coupled receptor (*GPCR*) mediated signaling, and ionic homeostasis in the infected livers. *KCNJ13,* which mediates potassium ion transmembrane transport, was upregulated. *TRPV4* (transient receptor potential cation channel subfamily V member 4), a calcium channel that regulates the production of pro-inflammatory chemokines, was upregulated in SLD-affected livers.

#### DNA repair and transport

Genes associated with genome maintenance and transport functions were downregulated, including *RAD51C* (homologous recombination), *SLC18A1* (serotonin uptake), and *SLC28A2* (nucleoside transport), indicating compromised cellular integrity in the terminal SLD.

#### Apoptotic regulation

Genes encoding key regulators of apoptosis, including *ITPRIP* (extrinsic apoptotic signaling via death domain receptors), *PLEKHO2* (macrophage apoptotic processes), and *C1QTNF1*, were significantly upregulated, consistent with the activation of programmed cell death pathways during hepatic damage.

### 3.5 Confirmation of RNA-seq Results with RT-qPCR

To validate the transcriptional profiles identified by RNA-seq, four representative genes (*IL-22, IL-6, RAD51C,* and *FBP2*) were analyzed using RT-qPCR. As shown in Fig 5, the expression trends observed by RT-qPCR were consistent with the RNA-seq data in liver samples, with both methods showing similar patterns of upregulation or downregulation. These results confirm the reliability of the RNA-seq expression profiles.

**Fig 5.**
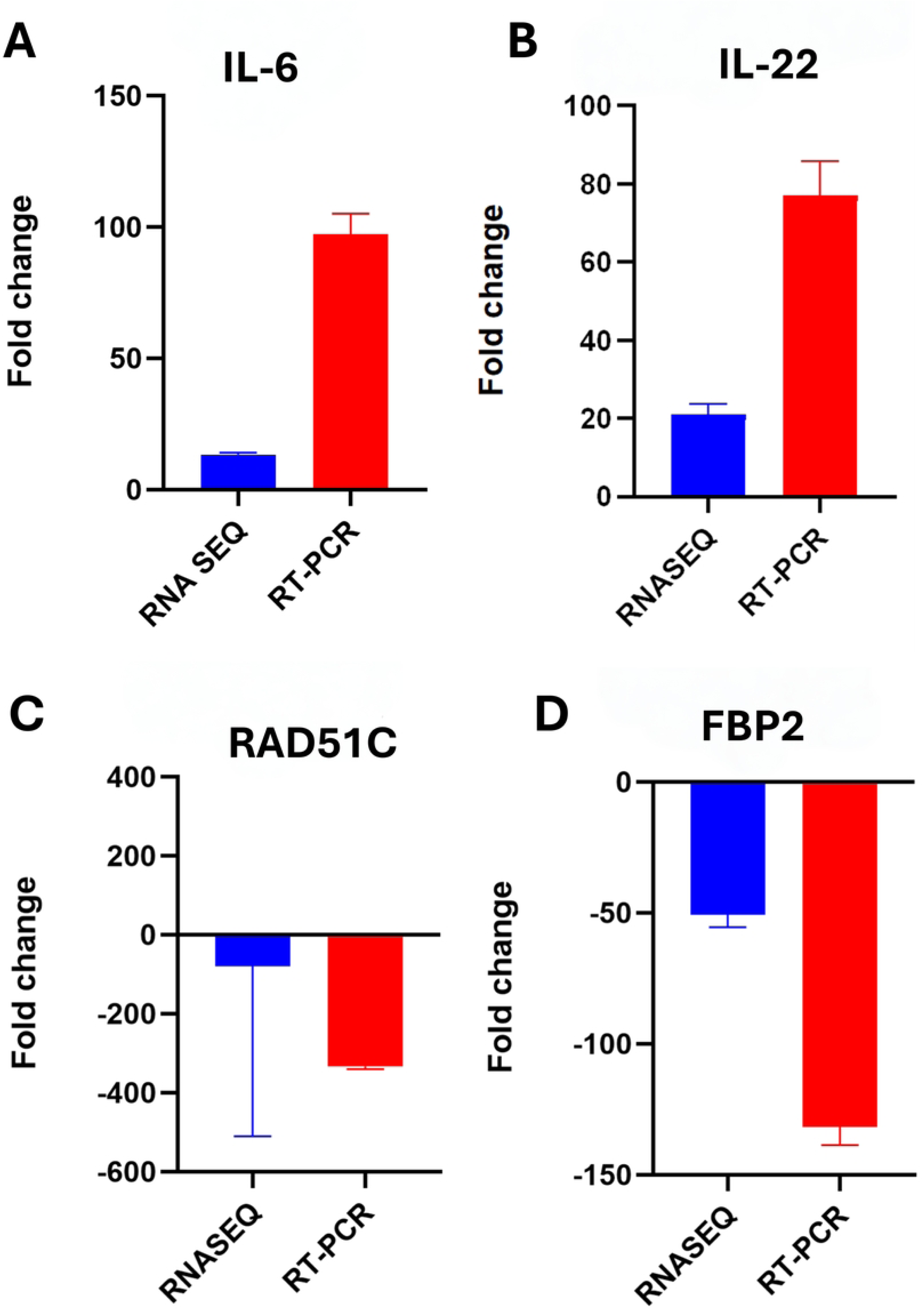
**Validation of RNA-seq data by RT-qPCR**. Relative expression levels of four representative chicken genes, *IL-6* (A), *IL-22* (B), *RAD51C* (C), and *FBP2* (D) in liver samples were compared between RNA-seq (blue bars) and RT-qPCR (red bars).

## Discussion

Transcriptome analysis of chicken liver affected by SLD revealed a complex host-pathogen relationship marked by significant immunological dysregulation, metabolic disruption, and cellular failure. These findings shed light on the molecular mechanisms underlying the severe pathology of SLD, distinguishing it from other *Campylobacter* infections that are often asymptomatic in chickens.

The pathway enrichment analysis revealed robust activation of immune-related processes, cytokine-mediated signaling, intestinal immune network for IgA production, MAPK signaling pathway, and cell surface receptor signaling pathway among the most significantly enriched pathways associated with upregulated genes. Upregulation of key cytokine genes, including *IL2RB, IL-21, IL-22, IL8L2, IL-6, IL17REL,* and *IL17RE*, reflects activation of distinct immune cell populations and a coordinated response to *C. hepaticus* invasion. IL-22, a member of the IL-10 cytokine family that directly targets hepatocytes to stimulate antimicrobial peptide production and pro-inflammatory cytokine release (21) was markedly upregulated in SLD-affected livers.

This upregulation reflects the host’s attempt to initiate a protective response, as IL-22 typically promotes hepatocyte survival and antimicrobial defense. However, IL-22 exhibits a dual role in inflammatory diseases, functioning as either protective or pathogenic depending on the context (22). In SLD, the persistence of hepatic damage despite robust IL-22 induction suggests that the protective effects of this cytokine are overwhelmed by the extensive cellular stress and tissue pathology caused by *C. hepaticus* infection. The IL-6 gene was also upregulated in infected liver tissue. Previous studies have consistently identified IL-6 as a key inflammatory mediator in both *C. jejuni* and *C. hepaticus* infections, where it serves as an immediate marker for inflammation alongside other proinflammatory cytokines marking the onset of inflammation (15). Acute-phase proteins (APPs), largely synthesized by hepatocytes in response to pro-inflammatory cytokines, are released after infection or inflammation. SAA is the major and most sensitive APP in chickens, and was upregulated in SLD-affected liver tissues, confirming a severe hepatic inflammatory response and extensive hepatocyte activation during *C. hepaticus* infection (23). IL-21 plays a dual immunomodulatory role by enhancing T cell activation while simultaneously blocking dendritic cell maturation, which may contribute to the dysregulated immune response observed in terminal SLD. The concurrent upregulation of the IL-21 gene alongside its ability to inhibit dendritic cell maturation may paradoxically impair antigen presentation and contribute to the unsuccessful pathogen clearance observed in fatal SLD cases (24). Additionally, the strong upregulation of *CXCL13* in SLD-affected liver tissues indicates activation of B cell recruitment pathways, as chicken CXCL13 variants function as the primary B cell-attracting chemokines through binding to CXCR5 receptors on B cells and T cell subsets. Similarly, the upregulation of *ACKR4* (atypical chemokine receptor 4) in *C. hepaticus*-infected liver tissues indicates activation of chemokine scavenging mechanisms. In chickens, scavenger receptors are known to play important roles in innate immune defenses and to stimulate dose-dependent inflammatory responses in macrophages (25). The upregulation of *LYGL* (lysozyme G-like) and avian β-defensin (*AVBD*) genes indicates activation of innate antimicrobial defense mechanisms in SLD-affected livers. Lysozyme G-like proteins degrade bacterial peptidoglycan cell walls, providing direct bactericidal activity. Avian β-defensins are antimicrobial peptides with potent activity against multiple bacterial species, including *Campylobacter* (26). Previous studies have demonstrated that *C.* jejuni infection induces dynamic expression of avian host defense peptides, with significant upregulation of multiple AVBD genes in infected chickens (27). The marked upregulation of *AVBD1* in *C. hepaticus*-infected livers suggests hepatocytes are mounting a vigorous antimicrobial response to bacterial challenge. However, the persistence of severe hepatic lesions and high mortality in SLD despite robust antimicrobial gene upregulation indicate that *C. hepaticus* may have evolved mechanisms to evade these defenses in the hepatic microenvironment . Conversely, the downregulation of NK and T cell activation (28) and multiple immune-regulatory loci reflect suppression of adaptive immune control mechanisms during terminal SLD. These findings suggest that *C. hepaticus* establishes an immunosuppressive microenvironment, enabling bacterial persistence despite the robust inflammatory response observed.

The pathway enrichment analysis of downregulated genes revealed profound suppression of core metabolic functions. The genes involved in oxidative phosphorylation, fatty acid metabolic processes, carbon metabolism, fatty acid degradation, iron ion binding, heme binding, Peroxisome Proliferator-Activated Receptor (PPAR) signaling pathway were significantly downregulated, reflecting a profound impairment of mitochondrial and cellular core metabolic processes. Collectively, these alterations suggest that *C. hepaticus* infection disrupts metabolic processes in hepatocytes, contributing to hepatic failure and the characteristic pathology observed in SLD. This pattern of metabolic suppression is consistent with findings from other avian bacterial liver infections (29) where metabolic genes are typically downregulated during the acute phase response. However, the extent of metabolic dysfunction in SLD exceeds what has been reported for other avian pathogens, with the downregulation of genes encoding key glucose metabolism genes *FBP2, ALDOB and HKDC1* indicating comprehensive disruption of glucose homeostasis (30). The suppression of lipid metabolism genes, including *APOB, APOA2, and PLIN1*, reflects severe disruption of hepatic lipid transport and storage functions (31). The downregulation of *OTC* (ornithine transcarbamylase gene) indicates a compromised ammonia detoxification capacity, which is particularly problematic in chickens, given their differential ammonia metabolism compared to mammals. The concurrent suppression of cytochrome P450 enzyme genes *CYP2C18* and *CYP2AC1* indicates severely compromised xenobiotic metabolism (32), while the downregulation of *ACAA1* reflects disrupted mitochondrial fatty acid β-oxidation and energy production. This metabolic and energy failure not only threatens hepatocyte survival but also compromises the liver’s role as the primary site for yolk precursor synthesis in laying hens. The suppression of fatty acid synthesis pathways is expected to decrease triglyceride and phospholipid production (33) resulting in reduced egg production and inferior egg quality, clinical observations consistent with field reports of SLD-affected flocks. *C.hepaticus* genome exhibits a reduced repertoire of genes associated with iron acquisition and metabolism (34), indicating adaptation to iron-rich host environments like the liver. In chickens, the liver functions as the primary site for iron storage and regulation, maintaining systemic iron balance. Infected livers exhibited significant downregulation of pathways associated with iron ion, heme, and tetrapyrrole binding pathways, suggesting a localized modulation of hepatic iron homeostasis, potentially increasing iron bioavailability within the hepatic microenvironment, thereby favoring bacterial persistence and exacerbating disease progression. Notably, our previous studies on bacterial transcriptomic analysis of *C. hepaticus* during infection LMH cell line demonstrated upregulation of the siderophore transport gene *ceuB* (35). When considered together, the host downregulation of iron-binding pathways and the bacterial upregulation of *ceuB* support a model in which *C. hepaticus* exploits infection-induced disruption of hepatic iron homeostasis to secure iron within the hepatic microenvironment, thereby promoting persistence and disease progression. The dramatic upregulation of *C1QTNF1* (C1q and TNF-related protein 1) and *ADGRD1* genes in SLD-infected liver tissues represents a critical finding with implications for both inflammatory responses and cellular stress adaptation. C1QTNF1 functions as both a complement-related immune effector and metabolic regulator, with its upregulation suggesting activation of complement-mediated inflammatory pathways consistent with the severe hepatic lesions characteristic of terminal-stage disease (36). Similarly, ADGRD1, although not well characterized in chickens, is an adhesion G protein-coupled receptor known in mammals to mediate hypoxia and inflammatory-signaling. This pronounced upregulation during *C. hepaticus* infection (37) suggesting that hepatocytes are undergoing substantial stress and attempting to cope with the combined metabolic and inflammatory burden imposed by infection.

## Conclusions

In summary, transcriptomic profiling of SLD-affected chicken livers revealed marked immune dysregulation accompanied by profound suppression of core metabolic pathways. This convergence of immune dysregulation, including inflammatory imbalance and metabolic disruption, offers mechanistic insight into the high mortality and egg-production losses observed in affected flocks. The identification of key cytokines, acute-phase proteins, and metabolic genes may uncover promising biomarkers for disease surveillance and therapeutic intervention.

Importantly, this study suggests adaptive strategies by which *C. hepaticus* overcomes limitations in iron acquisition within the hepatic microenvironment despite the reduced repertoire of genes associated with iron acquisition and metabolism in its genome. Together, these findings establish a molecular framework for developing targeted diagnostic tools and control strategies to reduce the economic and welfare burden of SLD in commercial layer operations.

## Supplementary Materials

S1 File: Primers used for RT-qPCR analysis; S2 File: Overall differential gene expression profile data control liver and infected liver; S3 File: GO and KEGG analysis list of upregulated genes ; S4 File: GO and KEGG analysis list of downregulated genes

## References

Arukha, A., Denagamage, T. N., Butcher, G., & Kariyawasam, S. (2021). Complete Genome Sequence of Campylobacter hepaticus Strain UF2019SK1, Isolated from a Commercial Layer Flock in the United States. 10.1016/j.vetmic

Chagneau, S., Gaucher, M. Lou, Fravalo, P., Thériault, W. P., & Thibodeau, A. (2023). Intestinal Colonization of Campylobacter jejuni and Its Hepatic Dissemination Are Associated with Local and Systemic Immune Responses in Broiler Chickens. Microorganisms, 11(7). 10.3390/microorganisms11071677

Chen, M., He, Y., Jia, Y., Wu, L., & Zhao, R. (2025). Liver transcriptome response to avian pathogenic Escherichia coli infection in broilers with corticosterone treatment. Poultry Science, 104(5). 10.1016/j.psj.2025.105020

Coble, D. J., Sandford, E. E., Ji, T., Abernathy, J., Fleming, D., Zhou, H., & Lamont, S. J. (2013). Impacts of Salmonella enteritidis infection on liver transcriptome in broilers. Genesis, 51(5), 357–364. 10.1002/dvg.22351

Courtice, J. M., Ahmad, T. B., Wei, C., Mahdi, L. K., Palmieri, C., Juma, S., Groves, P. J., Hancock, K., Korolik, V., Petrovsky, N., & Kotiw, M. (2023). Detection, characterization, and persistence of Campylobacter hepaticus, the cause of spotty liver disease in layer hens. Poultry Science, 102(7), 102462. 10.1016/J.PSJ.2022.102462

Courtice, J. M., Mahdi, L. K., Groves, P. J., & Kotiw, M. (2018). Spotty Liver Disease: A review of an ongoing challenge in commercial free-range egg production. Veterinary Microbiology, 227, 112–118. 10.1016/J.VETMIC.2018.08.004

Crawshaw, T. (2019). A review of the novel thermophilic Campylobacter, Campylobacter hepaticus, a pathogen of poultry. In Transboundary and Emerging Diseases (Vol. 66, Number 4, pp. 1481–1492). Blackwell Publishing Ltd. 10.1111/tbed.13229

Crawshaw, T. R., Chanter, J. I., Young, S. C. L., Cawthraw, S., Whatmore, A. M., Koylass, M. S., Vidal, A. B., Salguero, F. J., & Irvine, R. M. (2015). Isolation of a novel thermophilic Campylobacter from cases of spotty liver disease in laying hens and experimental reproduction of infection and microscopic pathology. Veterinary Microbiology, 179(3–4), 315–321. 10.1016/j.vetmic.2015.06.008

Cuperus, T., Coorens, M., van Dijk, A., & Haagsman, H. P. (2013). Avian host defense peptides. Developmental and Comparative Immunology, 41(3), 352–369. 10.1016/j.dci.2013.04.019

Eastwood, S., Wilson, T. B., Huang, J., Campbell, B. E., Scott, P. C., Moore, R. J., & Van, T. T. H. (2025). Immune responses and recovery from spotty liver disease in layer birds. Poultry Science, 104(8), 105351. 10.1016/J.PSJ.2025.105351

Edison, L. K., & Kariyawasam, S. (2025). From the Gut to the Brain: Transcriptomic Insights into Neonatal Meningitis Escherichia coli Across Diverse Host Niches. Pathogens, 14(5). 10.3390/pathogens14050485

Edison, L. K., Kudva, I. T., & Kariyawasam, S. (2023). Comparative Transcriptome Analysis of Shiga Toxin-Producing Escherichia coli O157:H7 on Bovine Rectoanal Junction Cells and Human Colonic Epithelial Cells during Initial Adherence. Microorganisms, 11(10). 10.3390/microorganisms11102562

Garcia, J. S., Byrd, J. A., & Wong, E. A. (2018). Expression of nutrient transporters and host defense peptides in Campylobacter challenged broilers. Poultry Science, 97(10), 3671–3680. 10.3382/ps/pey228

Gottapu, C., Sahin, O., K. Edison, L., Srednik, M. E., & Kariyawasam, S. (2025). Complete genome sequences of Campylobacter hepaticus strains USA1 and USA5 isolated from a commercial layer flock in the United States . Microbiology Resource Announcements, 14(5). 10.1128/mra.00919-24

Gregory, M., Klein, B., Sahin, O., & Girgis, G. (2018). Isolation and Characterization of Campylobacter hepaticus from Layer Chickens with Spotty Liver Disease in the United States. Avian Diseases, 62(1), 79–85. 10.1637/11752-092017-REG.1

Hananeh, W., & Ababneh, M. (2021). Spotty liver disease in Jordan: An emerging disease. Veterinarni Medicina, 66(3), 94–98. 10.17221/73/2020-VETMED

He, H., MacKinnon, K. M., Genovese, K. J., Nerren, J. R., Swaggerty, C. L., Nisbet, D. J., & Kogut, M. H. (2009). Chicken scavenger receptors and their ligand-induced cellular immune responses. Molecular Immunology, 46(11–12), 2218–2225. 10.1016/J.MOLIMM.2009.04.020

Kim, S., Faris, L., Cox, C. M., Sumners, L. H., Jenkins, M. C., Fetterer, R. H., Miska, K. B., & Dalloul, R. A. (2012). Molecular characterization and immunological roles of avian IL-22 and its soluble receptor IL-22 binding protein. Cytokine, 60(3), 815–827. 10.1016/J.CYTO.2012.08.005

Liu, X. ting, Lin, X., Mi, Y. ling, Zeng, W. dong, & Zhang, C. qiao. (2018). Age-related changes of yolk precursor formation in the liver of laying hens. Journal of Zhejiang University: Science B, 19(5), 390–399. 10.1631/jzus.B1700054

Moore, R. J., Scott, P. C., & Van, T. T. H. (2019). Spotlight on avian pathology: Campylobacter hepaticus, the cause of Spotty Liver Disease in layers. In Avian Pathology (Vol. 48, Number 4, pp. 285–287). Taylor and Francis Ltd. 10.1080/03079457.2019.1602247

Petrovska, L., Tang, Y., Jansen van Rensburg, M. J., Cawthraw, S., Nunez, J., Sheppard, S. K., Ellis, R. J., Whatmore, A. M., Crawshaw, T. R., & Irvine, R. M. (2017). Genome reduction for niche association in Campylobacter hepaticus, a cause of spotty liver disease in poultry. Frontiers in Cellular and Infection Microbiology, 7(AUG). 10.3389/fcimb.2017.00354

Pilkis, S. J., El-Maghrabi, M. R., & Claus, T. H. (1990). Fructose∼2,6-Bisphosphate in Control of Hepatic Gluconeogenesis From Metabolites to Molecular Genetics. In 582 DIABETES CARE (Vol. 13, Number 6). http://diabetesjournals.org/care/article-pdf/13/6/582/439382/13-6-582.pdf

*PROCEEDINGS OF THE SIXTIETH WESTERN POULTRY DISEASE CONFERENCE*. (n.d.).

Quinteros, J. A., Scott, P. C., Wilson, T. B., Anwar, A. M., Scott, T., Muralidharan, C., Van, T. T. H., & Moore, R. J. (2021). Isoquinoline alkaloids induce partial protection of laying hens from the impact of Campylobacter hepaticus (spotty liver disease) challenge. Poultry Science, 100(11), 101423. 10.1016/J.PSJ.2021.101423

Ren, J., Yang, L., Li, Q., Zhang, Q., Sun, C., Liu, X., & Yang, N. (2019). Global investigation of cytochrome P450 genes in the chicken genome. Genes, 10(8). 10.3390/genes10080617

Risatti, G. R., Varga, C., Hänninen, M.-L., & Hao Van, T. T. (n.d.). *Prevalence of Campylobacter hepaticus specific antibodies among commercial free-range layers in Australia*.

Rothwell, L., Hu, T., Wu, Z., & Kaiser, P. (2012). Chicken interleukin-21 is costimulatory for T cells and blocks maturation of dendritic cells. Developmental & Comparative Immunology, 36(2), 475–482. 10.1016/J.DCI.2011.08.013

Van, T. T. H., Phung, C., Anwar, A., Wilson, T. B., Scott, P. C., & Moore, R. J. (2023). Campylobacter bilis, the second novel Campylobacter species isolated from chickens with Spotty Liver Disease, can cause the disease. Veterinary Microbiology, 276, 109603. 10.1016/J.VETMIC.2022.109603

Woo, S. J., Kim, J., Lee, H. J., Lee, K. Y., Park, K. J., Kim, J. K., Kim, J. L., Park, B. C., Seo, M., & Han, J. Y. (2025). Conserved functional features of natural killer cell subsets in chicken, human, and murine immune systems. IScience, 28(8). 10.1016/j.isci.2025.113144

Yu, H., Yan, X., Chen, G., Li, R., Yang, Z., Liang, Z., Ye, L., Chen, Y., & Li, Y. (2024). Dynamic network biomarker C1QTNF1 regulates tumor formation at the tipping point of hepatocellular carcinoma. Biomolecules and Biomedicine, 24(4), 939–951. 10.17305/bb.2024.10103

Žáčková, S., Pávová, M., Trylčová, J., Chalupová, J., Priss, A., Lukšan, O., & Weber, J. (2024). Upregulation of mRNA Expression of ADGRD1/GPR133 and ADGRG7/GPR128 in SARS-CoV-2-Infected Lung Adenocarcinoma Calu-3 Cells. Cells, 13(10). 10.3390/cells13100791

Zaefarian, F., Abdollahi, M. R., Cowieson, A., & Ravindran, V. (2019). Avian liver: The forgotten organ. Animals, 9(2). 10.3390/ani9020063

Zenewicz, L. A. (2018). IL-22: There Is a Gap in Our Knowledge. ImmunoHorizons, 2(6), 198–207. 10.4049/immunohorizons.1800006

Zhai, G., Pang, Y., Zou, Y., Wang, X., Liu, J., Zhang, Q., Cao, Z., Wang, N., Li, H., & Wang, Y. (2023). Effects of PLIN1 Gene Knockout on the Proliferation, Apoptosis, Differentiation and Lipolysis of Chicken Preadipocytes. Animals, 13(1). 10.3390/ani13010092

